# Restricted amino acid diversity alters enzymatic phosphoryl-transfer catalysis

**DOI:** 10.64898/2026.02.16.706253

**Authors:** Sota Yagi, Subrata Dasgupta, Shusuke Tagami, Satoshi Akanuma

## Abstract

How amino acid composition shapes enzymatic function remains a central question in molecular evolution. Early proteins were likely composed of a limited set of amino acids, yet the catalytic properties of proteins under such compositional constraints are still poorly understood. Here we show that restricting amino acid repertoires can replace native enzymatic activity with an alternative phosphoryl-transfer reaction that disproportionates ADP into ATP and AMP. Using a reconstructed ancestral nucleoside diphosphate kinase (NDK) variant composed of a restricted amino acid set, we demonstrate this reaction, which is not observed in extant NDKs but is chemically analogous to that catalyzed by adenylate kinase. Although modest in catalytic rate, X-ray crystallography, molecular dynamics, and mutational analyses reveal a distinct active-site organization in which aspartate and arginine residues cooperatively coordinate Mg^2+^ and support phosphoryl transfer through a noncanonical mechanism. Lysine can substitute for arginine under these constraints while retaining activity. Together, these findings show that restricting amino acid diversity can remodel active sites and promote alternative phosphoryl-transfer reactions, illustrating how limitations in amino acid availability could influence catalytic functions in early enzyme evolution.

**Significant Statement:** The first enzymes arose on early Earth, where only a limited set of amino acids was likely available. Whether such limitations reduce catalytic efficiency or reshape enzyme function is still unclear. We reconstructed a model of an ancestral enzyme using a simplified amino acid set approximating those available on early Earth and found that restricting amino acid diversity replaced its original catalytic activity with an alternative function that produces ATP from two ADP molecules. Structural and mutational analyses indicate that this shift involves a reorganization of the active site. These results provide experimental evidence that limiting amino acid diversity can reshape catalytic function and may have influenced the emergence of catalytic activities during enzyme evolution.

## Introduction

Understanding how proteins first emerged is a central issue in elucidating the transition from prebiotic chemistry to biological evolution^1–3^. One of the key challenges in this context is determining how the building blocks of proteins were initially acquired^4,5^. While modern proteins are generally composed of 20 L-amino acids, and it is believed that the last universal common ancestor (LUCA) could already synthesize proteins using 20 types of amino acids^6–8^, the amino acid composition of proteins prior to LUCA remains unclear^9–11^. To our best knowledge, there is no experimental evidence indicating that all 20 of the current proteinogenic amino acids were available during the earliest stages of life. More likely, these amino acids were instead gradually incorporated through the evolution of biosynthetic pathways^12–14^. Indeed, this proposal is supported by several lines of evidence. For example, ten amino acids were synthesized in Miller’s spark discharge experiment^15–17^, eight were identified in the Murchison meteorite^18,19^, and recent analyses of samples from asteroid Ryugu detected 11 proteinogenic amino acids, ten of which overlapped with those from the Miller experiment^20^. These findings suggest that primitive proteins were synthesized using only a limited subset of amino acids that were available on the early Earth, prior to the emergence of endogenous biosynthetic pathways. Specifically, the set of ten amino acids synthesized in Miller’s spark discharge experiment are often referred to as “prebiotic amino acids,” because they plausibly represent the amino acid repertoire available on early Earth^12,21,22^. However, modern enzymes rely heavily on the chemical diversity of side chains for substrate binding, transition-state stabilization, and catalysis. This raises a fundamental question of how primitive polypeptides, composed of a limited set of amino acids, could have folded into tertiary structures that exhibited enzymatic functions. Resolving this question is essential for reconstructing the molecular steps that led from simple prebiotic chemistry to the complex metabolic networks found in all known life.

The plausibility of primitive protein synthesis using the prebiotic set of amino acids has been supported by multiple lines of evidence. The set of prebiotic amino acids have been shown to associate with self-assembled fatty acid membranes, which enhances membrane stability. The localized concentration of these amino acids on the membrane surfaces could have facilitated peptide synthesis^21^. Computational and theoretical studies have further predicted that hypothetical proteins composed of the ten prebiotic amino acids would exhibit folding preferences and structural characteristics similar to those of modern proteins^23^. Moreover, another study suggested that among all possible combinations of ten amino acids drawn from the twenty used in modern proteins, the prebiotic set is particularly favorable for forming robust protein backbones^14^. Collectively, these findings indicate that the prebiotic amino acid subset was not only abundant in the early Earth environment but also inherently suited to forming stable, protein-like structures.

Nucleoside diphosphate kinase (NDK) catalyzes the transfer of the γ-phosphate from a nucleoside triphosphate to a nucleoside diphosphate, thereby maintaining the intracellular balance of nucleoside triphosphates^24^. Previously, we performed ancestral sequence reconstruction on NDK to produce several ancestral NDKs that might have existed in the last common ancestors of Archaea and Bacteria^25, 26^. Next, we systematically restricted the amino acid composition of Arc1, one of the archaeal ancestral NDKs, while preserving its catalytic function^27^. This approach yielded simplified proteins composed of only 13 amino acid types that retained detectable enzymatic activity and were thermostable above 70°C. Notably, the majority of amino acids in these simplified proteins overlapped with the aforementioned prebiotic amino acids. Furthermore, when histidine, asparagine, and tyrosine, which are not listed as prebiotic amino acids, were removed from a 13-amino acid variant, its high thermal stability was retained, although its catalytic activity was completely lost.

In this study, we extended our previous work by designing new variants with distinct amino acid compositions. Interestingly, one variant that was constructed using a 12-amino-acid set consisting of prebiotic amino acids supplemented with lysine and arginine exhibited a noncanonical catalytic activity: the synthesis of ATP and AMP from two ADP molecules, distinct from the canonical NDK reaction. This unexpected gain-of-function highlights a fundamental insight; namely, constraining amino acid composition can result in the emergence of alternative catalytic functions. ATP synthesis is thought to be crucial in early bioenergetics^28^. Thus, the emergence of ATP synthesis activity may reflect the role of early proteins composed of a restricted set of amino acids in primitive energy metabolism. Our findings provide a compelling example that reduced chemical complexity may have shaped the functional landscape of early proteins during the origin and early evolution of life.

## Results

### A 12-amino acid ancestral NDK variant catalyzes noncanonical ATP synthesis

To investigate the relationship between amino acid composition and catalytic function in ancestral proteins, we first constructed two new variants of the reconstructed ancestral NDK Arc1. One of these variants, designated Arc1-16, was designed with a unique 16-amino acid alphabet that excluded the residues histidine, asparagine, tyrosine and cysteine. Histidine, asparagine and tyrosine residues are thought to play important roles in the catalytic activity of Arc1^27^, and cysteine is a residue not present in original Arc1 (Table 1; Fig. S1; Data S1). In addition, we constructed a 12-amino acid variant, Arc1-12KR, which was composed of the ten prebiotic amino acids supplemented with two basic residues, lysine and arginine (Table 1; Fig. S1; Data S1). Note, the N-terminal Met was excluded from the compositional definition.

**Table 1.** Specific activities at 60 °C, oligomeric structure and amino acid composition of Arc1 and its reduced amino acid set variants.

|  | Arc1 | Arc1-13FKR | Arc1-13KMR | Arc1-12KR | Arc1-11K |
| --- | --- | --- | --- | --- | --- |
| Specific activity (canonical reaction) (U/mg) | 1900 ± 200 | < 0.10<sup>a</sup> | < 0.10<sup>a</sup> | n/a<sup>b</sup> | n/a<sup>b</sup> |
| Specific activity (ADP disproportionation) (U/mg) | < 0.10<sup>a</sup> | < 0.10<sup>a</sup> | < 0.10<sup>a</sup> | 5.0 ± 0.3 | 2.5 ± 0.0 |
| Oligomeric structure | Hexamer | Hexamer | Hexamer | Dimer | Dimer |
| Ala | 11 | 12 | 12 | 12 | 12 |
| Asp | 6 | 9 | 9 | 9 | 9 |
| Glu | 16 | 16 | 16 | 16 | 16 |
| Gly | 11 | 11 | 11 | 11 | 11 |
| Ile | 10 | 10 | 10 | 10 | 10 |
| Leu | 7 | 19 | 19 | 26 | 26 |
| Pro | 6 | 6 | 6 | 6 | 6 |
| Ser | 8 | 8 | 8 | 8 | 8 |
| Thr | 4 | 4 | 4 | 4 | 4 |
| Val | 14 | 14 | 14 | 14 | 14 |
| Lys | 11 | 11 | 11 | 11 | 22 |
| Arg | 10 | 11 | 11 | 11 | 0 |
| Phe | 7 | 7 | 0 | 0 | 0 |
| Met<sup>c</sup> | 7 | 0 | 7 | 0 | 0 |
| His | 3 | 0 | 0 | 0 | 0 |
| Asn | 3 | 0 | 0 | 0 | 0 |
| Gln | 1 | 0 | 0 | 0 | 0 |
| Trp | 1 | 0 | 0 | 0 | 0 |
| Tyr | 2 | 0 | 0 | 0 | 0 |
| Cys | 0 | 0 | 0 | 0 | 0 |

<sup>a</sup> The catalytic activities were below the limit of detection.
<sup>b</sup> The assay used does not distinguish ATP production via the canonical reaction from that via ADP disproportionation.
<sup>c</sup> The N-terminal residue was not considered.

Unless otherwise noted, all protein variants were expressed with an N-terminal His₆-tag. For Arc1-12KR, we confirmed that the His-tag did not affect folding or enzymatic activity by comparing the tagged and tag-free forms using circular dichroism (CD) spectroscopy, and ATP-synthesizing assays (Fig. S2). In addition, the constructs lacking aromatic amino acids were appended with a Gly-Gly-Trp tag at the C-terminus to enable spectroscopic detection of these proteins.

NDK catalyzes the transfer of the γ-phosphate from a nucleoside triphosphate to a nucleoside diphosphate (Fig. 1A). To assess whether the new variants retained this canonical activity, enzyme assays were performed in which ATP production was monitored using GTP and ADP as substrates. Because both variants lacked histidine, a residue shown to be essential for catalysis, none were expected to retain the original enzymatic activity. Consistent with this expectation, Arc1-16 did not display detectable catalytic activity under these conditions (Fig. 1C; Data S2). Intriguingly, however, ATP synthesis activity was detected for Arc1-12KR (Fig. 1C inset). More surprisingly, ATP formation was also observed when GTP, one of the substrates, was omitted from the reaction mixture (Fig. 1D; Table 1; Data S2). This result prompted us to hypothesize that Arc1-12KR was catalyzing an alternative reaction, namely, the formation of ATP and AMP from two molecules of ADP (Fig. 1B). To test this hypothesis, we analyzed the reaction products using high-performance liquid chromatography (HPLC). Prior to incubation, only a single peak corresponding to ADP was observed in the reaction mixture (Fig. 1E; Fig. S3). After incubation with Arc1-12KR at 60 °C for 10 min, two additional peaks emerged, corresponding to ATP and AMP (Fig. 1E; Fig. S3). These results confirmed that Arc1-12KR catalyzes the formation of ATP and AMP from two molecules of ADP.

**Fig. 1.**
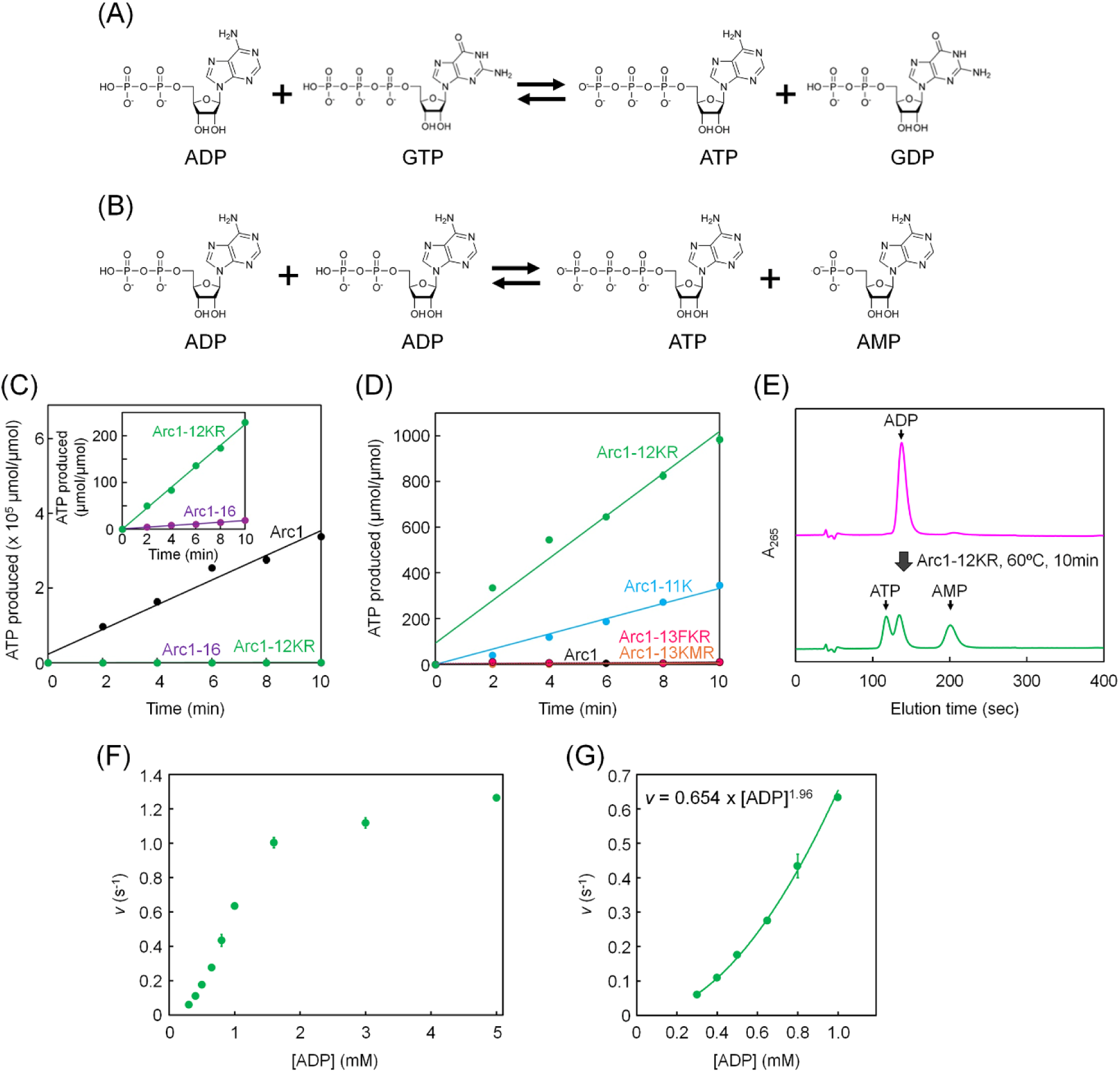
Reaction schemes and catalytic activities of canonical NDKs and reduced amino acid set variants. **(A)** Canonical NDK-catalyzing reaction for the transfer of γ-phosphate of a nucleoside triphosphate (e.g., GTP) to a nucleoside diphosphate (e.g., ADP). **(B)** ADP disproportionation reaction catalyzed by Arc1-12KR and its variants, producing ATP and AMP from two molecules of ADP. **(C)** Time-course of ATP formation catalyzed by Arc1, Arc1-16, and Arc1-12KR using ADP and GTP as substrates, corresponding to the canonical NDK-catalyzing reaction shown in panel (A). The inset shows an expanded view highlighting the activities of Arc1-16 and Arc1-12KR. **(D)** Time-course of ATP production from ADP as the sole substrate by Arc1 and its reduced amino acid set variants. **(E)** HPLC analysis of reaction products obtained after incubating ADP with Arc1-12KR at 60 °C for 10 min, demonstrating the formation of ATP and AMP as predicted by the disproportionation reaction shown in panel (B). **(F)** Initial reaction velocity (*v*) of ATP formation as a function of ADP concentration for the ADP disproportionation reaction catalyzed by Arc1-12KR (0.3–5.0 mM ADP). **(G)** Expanded view of the low ADP concentration range (0.3–1.0 mM) shown in panel (F). Power-law fitting yielded an exponent of approximately 1.96. This result is consistent with an apparent second-order dependence on ADP concentration.

To examine whether this unexpected activity was specific to Arc1-12KR, we tested the original Arc1 and two 13-amino acid-alphabet variants, Arc1-13FKR and Arc1-13KMR, in which all phenylalanine residues or all methionine residues present in the original Arc1 sequence were restored to the Arc1-12KR scaffold, respectively (Table 1; Fig. S1; Data S1). When the activity was measured with ADP as the only substrate, no detectable ATP production was observed above background levels for these constructs (Fig. 1D; Table 1; Data S2), indicating that the ability to catalyze ADP disproportionation is unique to Arc1-12KR under the conditions used in this study.

Next, we examined the rate of ATP formation as a function of ADP concentration (Fig. 1F; Data S2). In the low ADP concentration range (0.3–1.0 mM), the reaction rate determined from the linear phase of ATP formation exhibited an approximately second-order dependence on ADP concentration (Fig. 1G). This nonlinear relationship cannot be readily explained by a canonical ping-pong mechanism and is more consistent with a mechanism involving the direct transfer of a phosphate between two ADP molecules.

We also examined whether Arc1-12KR is able to catalyze the reverse reaction. Upon incubating the enzyme with ATP and AMP under the standard assay conditions (2 mM Mg^2+^), no detectable formation of ADP was observed (Fig. S4A). However, increasing the Mg^2+^ concentration to 10 mM resulted in detectable ADP formation from ATP and AMP (Fig. S4B), suggesting that the reverse reaction depends on Mg^2+^ availability relative to nucleotide concentration. Furthermore, when GDP was used in place of ADP, no reaction products were detected (Fig. S4C), indicating that the observed activity is specific to adenylates and does not reflect a generalized NDK activity.

To assess the oligomeric states of Arc1-12KR and its related variants, we performed analytical gel filtration (Fig. S5). Arc1-13FKR and Arc1-13KMR eluted at volumes corresponding to their respective hexameric assemblies, similar to the oligomeric state of Arc1 and many extant NDKs^29–31^. In contrast, Arc1-12KR eluted as a single peak corresponding to a dimer. Notably, Arc1-12KR, which uniquely exhibited the altered catalytic activity, existed predominantly as a dimer, whereas the hexameric variants did not display this activity. Structural analysis indicates that the interfaces involved in hexamer formation do not directly contribute to the active site. Therefore, while the change in oligomeric state correlates with the altered activity, a causal relationship between oligomerization and functional divergence cannot be established.

### Dissecting the roles of arginine and lysine in noncanonical ATP synthesis

To further dissect the contribution of basic amino acids in the newly observed catalytic activity of Arc1-12KR, we designed three additional variants: Arc1-11K, in which all arginine residues were replaced with lysine (Table 1; Fig. S1; Data S1); Arc1-11R, in which all lysine residues were replaced with arginine (Fig. S1; Data S1); and Arc1-10, in which both lysine and arginine residues were replaced with prebiotic amino acids selected from the remaining ten used in Arc1-12KR (Fig. S1; Data S1). The corresponding genes were codon-optimized for expression in *Escherichia coli* and artificially synthesized. Despite codon optimization, no expression was detected for either Arc1-11R or Arc1-10 in *E. coli* under our standard expression conditions, suggesting that these variants did not fold into an adequate tertiary structure and were readily degraded. Due to this poor expression, Arc1-11R and Arc1-10 were not purified and structurally characterized. In contrast, Arc1-11K was expressed at high levels and the recombinant protein successfully purified for further analysis.

Gel filtration chromatography revealed that Arc1-11K, like Arc1-12KR, exists predominantly as a dimer (Table 1; Fig. S5). Far-UV circular dichroism spectroscopy demonstrated that Arc1-11K exhibits a secondary structure content highly reminiscent of Arc1-12KR (Fig. S6). These findings indicated that the lysine residues could be substituted for arginine without disrupting the overall protein fold. Catalytic activity assays using ADP as the sole substrate showed that Arc1-11K retained the ability to catalyze the conversion of two ADP molecules into ATP and AMP (Fig. 1D; Table 1; Data S2). However, the rate of reaction was approximately half that of Arc1-12KR, suggesting arginine contributes more effectively than lysine to this noncanonical catalytic function.

### Structural basis of histidine-independent ATP synthesis mediated by Arc1-12KR

To gain insight into the structural basis of the unexpected catalytic activity observed in Arc1-12KR, we determined the crystal structure of the apo form of this variant at 3.3 Å resolution by X-ray crystallography (Table S1). The asymmetric unit contained six independent dimeric assemblies (Fig. S7A), consistent with gel filtration data indicating that the protein predominantly exists as a dimer in solution. Superposition of the six dimers revealed notable conformational heterogeneity within a loop region (residues 49–59) located adjacent to the putative active site (Figs. S7B and S7C). This heterogeneity suggests inherent flexibility at the loop region, which may be relevant to the catalytic mechanism. Except for this flexible motif, the six protomers exhibited highly conserved backbone conformations, indicating that the overall fold is well maintained despite the reduced amino acid alphabet. Next, the crystal structure of Arc1-12KR was compared with that of the ancestral Arc1 protein (Fig. 2A). The two structures aligned closely, with the majority of backbone atoms showing excellent overlap. However significant deviation was observed in the loop formed by residues 49-59, which is located near the active-site and likely involved in substrate recognition and positioning (Fig. S8).

**Fig. 2.**
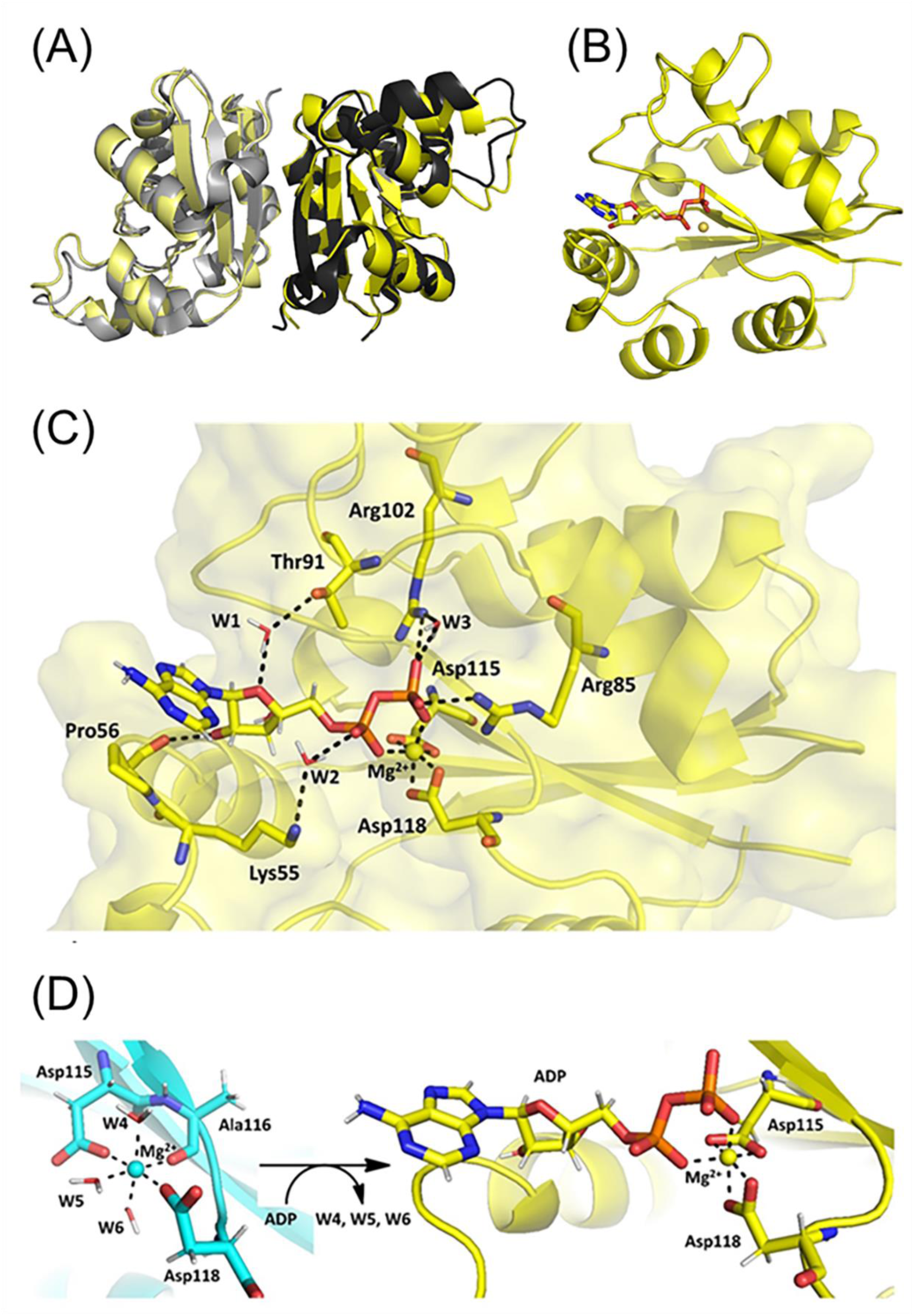
Crystal structure of Arc1-12KR and its MD-generated ADP–Mg^2+^ complex model. **(A)** Superposition of the Arc1-12KR dimer (yellow) with the crystal structure of Arc1 (black and gray, PDB code; 3VVT). **(B)** Model of the Arc1-12KR–ADP–Mg^2+^ complex generated by MD simulation. **(C)** Direct and water-mediated hydrogen bonds involving W1, W2, and W3 stabilize ADP binding in the catalytic pocket of the Arc1-12KR–ADP–Mg^2+^ complex model. **(D)** Comparison of Mg^2+^ coordination in the apo and holo forms. In the apo form (cyan), the Mg^2+^ ion is coordinated by Asp115, the main-chain carbonyl oxygen of Ala116, Asp118, and three conserved water molecules (W4, W5, and W6). In the holo form (yellow), these water molecules are displaced by ADP, and the Mg^2+^ ion is coordinated by Asp115, Asp118, and the α- and β-phosphate groups of ADP.

To directly visualize substrate binding, we attempted co-crystallization of Arc1-12KR with ADP. However, electron density maps revealed no interpretable density corresponding to ADP in any of the datasets obtained. Instead, a structural model of the Arc1-12KR–Mg^2+^–ADP complex was generated using molecular dynamics (MD) simulations (Fig. S9). Starting from the most “opened” apo conformation observed among the six crystallographically independent dimers, ADP molecules were docked into the putative active site. In addition, MD simulations were performed for the apo form of Arc1-12KR to provide a reference for comparison with the ADP-bound model.

The modeled complex structure revealed that ADP is located in the canonical binding pocket, with its phosphate groups coordinated by a central Mg^2+^ ion (Figs. 2B and 2C). In the modeled active site, the side chain guanidinium groups of Arg85 and Arg102 directly interact with the β-phosphate group of ADP through hydrogen bonds, indicating that these two arginine residues play important roles in catalysis. During the MD simulation, three water molecules (W1, W2, and W3) exhibited nearly 100% occupancy and contributed to ADP binding through hydrogen-bond networks (Fig. 2D). The hydroxyl group of Thr91 forms a hydrogen bond with ADP via W1. The side chain amino group of Lys55 forms a hydrogen bond with the α-phosphate of ADP via W2. In addition, the side chain guanidinium group of Arg102 interacts with ADP through W3. In the apo form, the Mg^2+^ ion adopts a distorted octahedral coordination geometry, involving the side chain carboxylate groups of Asp115 and Asp118, the main-chain carbonyl oxygen of Ala116, and one water molecule (W5) in the equatorial plane, while two additional water molecules (W4 and W6) occupy the axial positions. In the ADP-bound form, these water molecules are displaced by ADP, and the Mg^2+^ ion is coordinated by the side chain carboxylate groups of Asp115 and Asp118 as well as the α- and β-phosphate groups of ADP.

The active sites of the modeled Arc1-12KR and the *Pyrococcus horikoshii* OT3 NDK-ADP-Mg^2+^ complex (PDB ID: 2DY9) are compared in Fig. 3 and superimposed in Fig. S10. Several critical differences between the two active sites were observed. The bound ADP adopts different positions in the active site and, therefore, the phosphorus atom of the β-phosphate is shifted by ∼1.3 Å, and the position of the coordinated Mg^2+^ ion differs by ∼2.2 Å between the two structures. Residue 115, which is a catalytically essential histidine in all known NDKs, is replaced by an aspartate in Arc1-12KR. In canonical NDKs, including that from *P. horikoshii*, the conserved histidine residue functions as a transient phosphoryl acceptor during catalysis. In Arc1-12KR, however, Asp115 is positioned in close proximity to the catalytic Mg^2+^ ion, suggesting that it contributes primarily to metal coordination rather than directly participating in phosphoryl transfer. The carboxylate moiety of Asp118 in Arc1-12KR is also positioned close to the Mg^2+^ ion, whereas the corresponding aspartate residue in *P. horikoshii* NDK is oriented away from the active site. In contrast, Arg85 and Arg102 in Arc1-12KR and the corresponding arginine residues in *P. horikoshii* NDK are similarly positioned and oriented in the active site. This conservation of relative spatial positioning and orientation suggests that these arginine side chains play comparable roles in phosphate binding. The structural analyses suggest that Arc1-12KR adopts a distinct catalytic strategy primarily derived from alterations to acidic residues and metal coordination, while preserving the canonical phosphate-binding arginine residues. The active site architecture in Arc1-12KR reflects adaptation resulting from a restricted amino acid alphabet. These findings demonstrate how simplified sequence information can generate functional enzymatic folds with novel mechanistic features.

**Fig. 3.**
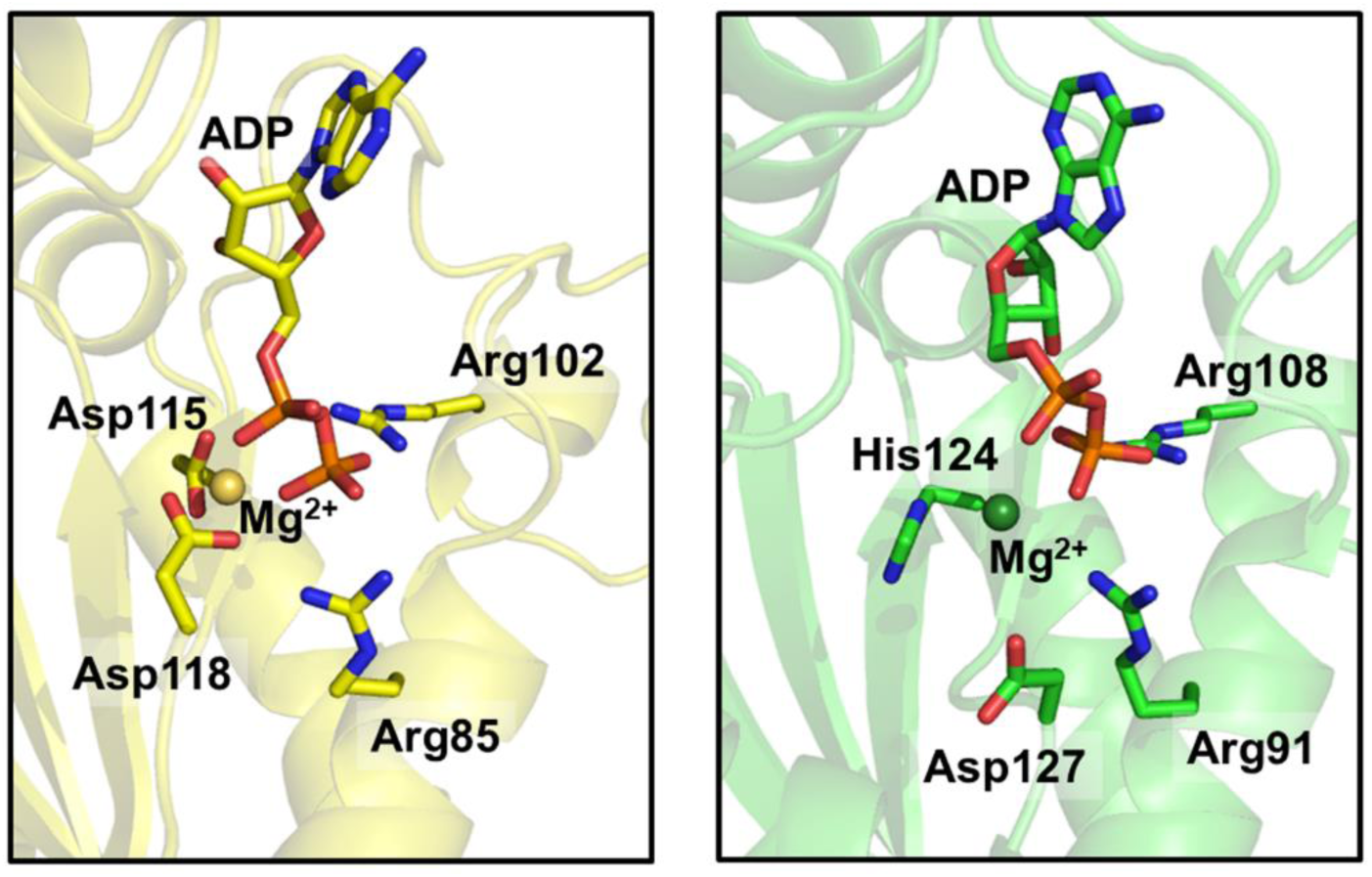
Comparison of the ADP-binding pockets of the Arc1-12KR-ADP-Mg^2+^ complex model (left) and the crystal structure of *P. horikoshii* NDK bound to ADP and Mg^2+^ (right). The side chains of the four key residues in each structure are also shown.

### Mutational analysis identifies residues critical for noncanonical ATP synthesis

Next, we aimed to identify crucial residues for the newly acquired ATP synthesis activity. Mutants of four candidate residues suggested from analysis of the crystal structure of Arc1-12KR and its MD-generated ADP-bound model were generated. Asp115 and Asp118, both oriented toward the catalytic Mg^2+^ ion, were individually substituted with alanine. The resultant D115A and D118A mutants retained 60% and 68% of the original ATP synthesizing activity, respectively (Table 2; Fig. S11; Data S2). However, simultaneous substitution of both residues drastically reduced this activity to ∼6% (Table 2; Fig. S11; Data S2), suggesting that the presence of at least one negatively charged side chain is important for coordinating Mg^2+^ and for electrostatic contributions to ADP binding.

**Table 2.** Specific ATP-synthesizing activity via ADP disproportionation for Arc1-12KR and its variants at 60 °C.

| Mutants | Specific activity (U/mg) |
| --- | --- |
| Arc1-12KR | $5.0 \pm 0.3$ |
| D115A | $3.0 \pm 0.2$ |
| D118A | $3.4 \pm 0.4$ |
| D115A/D118A | $0.30 \pm 0.01$ |
| R85M | $0.67 \pm 0.04$ |
| R102M | $2.4 \pm 0.1$ |
| R85M/R102M | $0.21 \pm 0.01$ |

Arg85 and Arg102, predicted to interact with the phosphate moieties of ADP, were individually replaced with methionine. R102M maintained approximately 50% of the original ATP synthesizing activity, but R85M resulted in a significant reduction in activity to ∼13% (Table 2; Fig. S11; Data S2). The double mutant lacking both arginine residues displayed only ∼4% activity (Table 2; Fig. S11; Data S2), indicating the cooperative role of Arg85 and Arg105 in positioning the substrate and stabilizing the transition state. Notably, Arg85 appears to play a more central role than Arg102 in this variant, despite the fact that, in extant NDKs, both Arg85 and Arg102 are identified as catalytically important residues^32^.

Intriguingly, the variant Arc1-11K, in which all arginine residues including Arg85 and Arg102 were replaced with lysine, retained 50% of the ATP-synthesizing activity of Arc1-12KR (Fig. 1D; Table 1; Data S2). Therefore, the catalytic roles of Arg85 and Arg102 can be compensated by lysine, although with reduced efficiency. These findings likely reflect a compensatory redistribution of catalytic roles that emerged under the constraints of being restricted to a prebiotically plausible set of amino acids.

## Discussion

In this study, we show that ATP-synthesizing activity emerges in the reconstructed kinase-like enzymes Arc1-12KR and Arc1-11K even though they are built only from prebiotic and basic amino acids. As such, these enzymes operate under severe compositional constraints. The reaction converting two molecules of ADP into ATP and AMP is, among known enzymes, catalyzed by adenylate kinases^33^, but it has not been observed in extant NDKs or in the reconstructed ancestral NDK Arc1. Thus, within the NDK family, this ATP-synthesizing activity represents a highly unusual and genuinely novel function. From an evolutionary perspective, this activity may reflect a primitive mode of phosphoryl transfer that predates the appearance of histidine-containing active sites. A restricted amino-acid alphabet lacking histidine, asparagine, tyrosine and other late-emerging residues^34^ would have constrained conventional catalytic strategies and forced a repurposing of residue side chains such as arginine and aspartate for substrate binding, charge compensation and possibly phosphate transfer. Our results therefore indicate that changes in biochemical function can arise not only from increasing molecular complexity but also from the adaptive exploitation of physicochemical constraints.

The ability of Arc1-12KR to catalyze a reaction that is absent from its modern homologs suggests that structural plasticity and chemical adaptability may have played key roles in the functional diversification of primordial enzymes. Rather than representing an entirely novel catalytic activity, the observed enzymatic function may instead reveal a latent capacity that was accessible on early protein scaffolds but was subsequently lost as more efficient and specialized mechanisms evolved. Our findings demonstrate that compositional simplification of biocatalysts can unmask alternative activities that are no longer retained by modern, fully optimized enzymes.

Consistent with this notion, Arc1-12KR and Arc1-11K catalyze the same overall reaction as modern adenylate kinases despite having no detectable homology and entirely distinct tertiary structures. This functional convergence implies that phosphoryl transfer reactions may have arisen independently multiple times during early evolution from nonhomologous protein scaffolds. The canonical NDK reaction, which involves transfer of the γ-phosphate from NTP to NDP, is nearly universal across extant life. By contrast, the ADP disproportionation reaction characterized here may reflect an older stage of molecular evolution. In environments where NTPs were scarce or absent, a mechanism that uses ADP as both phosphate donor and acceptor could have provided a plausible route for transient ATP formation and energy recycling in prebiotic systems. As illustrated in Fig. 4, we envisage a prebiotic world in which kinase-like proteins resembling Arc1-12KR/Arc1-11K with a limited amino acid repertoire generated ATP via ADP disproportionation. With the subsequent expansion of the amino acid alphabet, this chemistry was later superseded by modern NDKs, adenylate kinases and rotary ATPases.

**Fig. 4.**
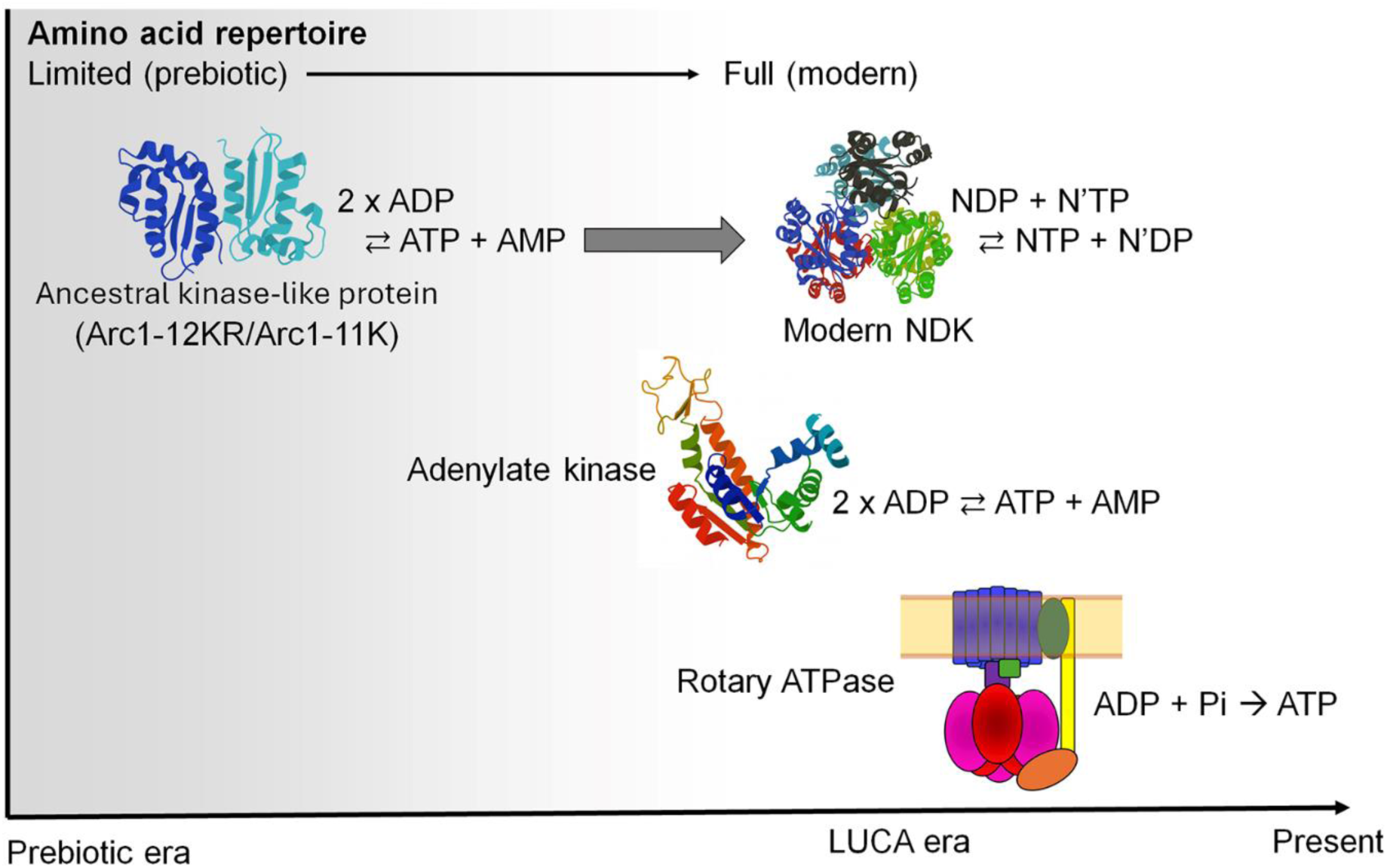
Conceptual model illustrating a possible evolutionary trajectory of ATP-generating enzymes. Under prebiotic conditions with a limited amino acid repertoire, kinase-like proteins resembling Arc1-12KR or Arc1-11K may have generated ATP through ADP disproportionation (2ADP → ATP + AMP). As the amino acid alphabet expanded toward the LUCA era, this chemistry was likely superseded by the emergence of modern NDKs, which catalyze phosphoryl transfer between NTPs and NDPs, and by adenylate kinases, which catalyze the reversible reaction 2ADP ⇌ ATP + AMP. Ultimately, the evolution of rotary ATP synthases enabled efficient ATP production from ADP and inorganic phosphate.

Although the ADP disproportionation reaction is thermodynamically close to neutral, its evolutionary and metabolic implications could be substantial. Local or transient increases in ATP concentration would have facilitated downstream biosynthetic reactions and early nucleotide-dependent processes^28^. At the same time, the co-produced AMP could have contributed to nucleotide signaling functions in primitive biochemical networks^33^. In proto-cellular environments, such reactions, when combined with compartmentalization or ion gradients, may have helped to create and sustain non-equilibrium states, thereby supporting the emergence of proto-metabolic organization before dedicated ATP synthase systems evolved^35,36^.

Canonical NDKs catalyze γ-phosphate transfer from NTP to NDP via a ping-pong mechanism involving a transient phosphohistidine intermediate^37^. This mechanism depends on a conserved histidine residue (His115 in Arc1), which serves as the nucleophile and is transiently phosphorylated during catalysis. By contrast, Arc1-12KR entirely lacks histidine and catalyzes disproportionation between two ADP molecules to yield ATP and AMP. Although aspartate residues have been reported to form transient phosphoaspartate intermediates in other enzymatic contexts^38,39^, mutational analyses indicate that neither Asp115 nor the nearby Asp118 is essential for ATP synthesis in Arc1-12KR, as alanine substitutions at these positions retain substantial activity (Table 2). Indeed, even the simultaneous substitution of both residues results in reduced but detectable catalytic activity. These observations argue against the formation of a covalent phosphoaspartate intermediate in Arc1-12KR and instead are consistent with a histidine-independent phosphoryl-transfer mechanism distinct from that of canonical NDKs. This interpretation is further supported by kinetic analyses showing that, at low ADP concentrations, the rate of ATP formation exhibits an approximately second-order dependence on ADP concentration (Fig. 1G). This kinetic data cannot be readily explained by a ping-pong mechanism and is more consistent with direct transfer of a phosphate moiety between two molecules of ADP.

Under the conditions used in this study (2 mM Mg^2+^), the ADP disproportionation reaction approached saturation at ∼1.6 mM ADP (Fig. 1F), and no detectable reverse reaction (ATP + AMP → 2 ADP) was observed (Fig. S4A). However, increasing the Mg^2+^ concentration to 10 mM resulted in detectable ADP formation from ATP and AMP (Fig. S4B), indicating that the apparent irreversibility reflects Mg^2+^ availability relative to nucleotide concentration. Given that nucleotide substrates are likely to enter the active site as Mg–ADP complexes, depletion of free Mg^2+^ at high ADP concentrations could account for the apparent saturation behavior. Moreover, preferential coordination of Mg^2+^ by ATP may reduce formation of Mg–AMP complexes under Mg^2+^-limiting conditions, thereby suppressing the reverse reaction.

Our structural and mutational analyses suggest that the noncanonical activity relies on cooperative interactions among Mg^2+^-coordinating aspartate and phosphate-binding arginine residues. Together, these residues may partially substitute for the catalytic role normally played by histidine in canonical NDKs. This occurs by organizing the Mg^2+^-ADP complex, stabilizing the negative charge on the phosphate groups, and facilitating phosphoryl transfer, which does not involve a covalent intermediate. These observations indicate that, when the amino acid alphabet is constrained, alternative catalytic mechanisms can emerge that diverge from those employed by modern enzymes.

The variant completely lacking basic amino acids (Arc1-10) failed to fold into a stable structure, highlighting a paradox; namely, although basic residues were probably rare under prebiotic conditions^40^, they seem to be essential for proper protein folding and activity. Previous work has shown that primitive proteins can be stabilized in high-salt environments^22^, and that proteins devoid of basic residues can interact with RNA via metal-ion mediation^41^. Future experiments should clarify how environmental factors, particularly the availability of mono and divalent cations, might have compensated for the absence of basic residues and enabled enzymatic activity in the early Earth environment.

However, we cannot rule out the possibility that lysine, arginine, or both these amino acids were in fact synthesized abiotically and already present in early protein synthesis^42,43^. Tawfik and co-workers proposed that the first nucleic acid-binding proteins may have arisen from short, simple sequences containing ornithine, which is not used in extant proteins^44^. By contrast, Thoma and Powner showed that ornithine and diaminobutyric acid rapidly cyclize intramolecularly, blocking further peptide extension, whereas the shortest basic α-amino acid diaminopropionic acid supports peptide formation and may therefore have substituted for or complemented lysine and arginine during early evolution^45^.

In conclusion, our work demonstrates that changes in enzyme function can arise not only through diversification and increasing structural complexity, but also through constraint and simplification of amino acid composition. By experimentally reconstructing conditions that approximate early evolution, we show that latent or lost catalytic potentials can be reactivated under restricted amino acid alphabets. Arc1-12KR and Arc1-11K may therefore exemplify how functional innovation, or reversion to ancient catalytic modes, can emerge when proteins are forced to operate within a limited repertoire of building blocks. The findings of this study support the view that the first enzymatic activities exploited prebiotic constraints to bridge the gap between nonenzymatic chemistry and genetically encoded biocatalysis, illuminating a plausible route toward the emergence of modern energy metabolism.

## Methods

### Design of Simplified NDK Sequences

To design amino acid sequences for the simplified NDK variants, residues targeted for elimination were substituted with the most frequent amino acids found at the corresponding positions in a multiple sequence alignment of 309 extant NDKs, excluding the residues chosen for removal (Data S3). For positions that were completely conserved, chemically similar amino acids were used instead. The N-terminal methionine residue as well as the tag sequences appended to the N- or C-termini were not included in these considerations.

### Expression and purification of simplified proteins

The simplified amino acid sequences were reverse-translated into nucleotide sequences optimized for *E. coli* codon usage. To enable spectroscopic detection of the expressed proteins, especially in the absence of aromatic residues such as tryptophan and tyrosine, all constructs were designed to include a Gly-Gly-Trp tripeptide tag at the C-terminus. In addition, we prepared both His-tagged and untagged forms of Arc1-12KR to assess the potential effect of the N-terminal His₆-tag on protein structure and activity (Fig. S2). The genes were synthesized by Eurofins Genomics (Tokyo, Japan) and cloned into the NdeI–BamHI restriction recognition sites of either pET15b (N-terminal His₆-tagged form) or pET23a(+) (untagged form) vectors (Merck, Tokyo, Japan). The resulting expression plasmids were used to transform *E. coli* Rosetta 2(DE3) cells (Merck), which were plated on Luria–Bertani (LB) agar containing 150 μg/mL ampicillin and incubated at 37 °C overnight. Several colonies were then inoculated into 200 mL of LB medium supplemented with ampicillin (150 μg/mL) and grown overnight at 37 °C. Protein overexpression was induced using the Overnight Express Autoinduction System 1 (Merck).

The cultured cells were harvested by centrifugation (5,000 × g, 10 min, 4 °C) and then disrupted by sonication. Lysates were heat-treated at 60, 65, or 70 °C for 20 min to precipitate host *E. coli* proteins. After centrifugation (20,000 × g, 30 min, 4 °C), His₆-tagged proteins were purified using a HisTrap HP column followed by gel filtration using a Superdex 200 Increase 10/300 GL column (GE Healthcare Japan, Tokyo, Japan). Untagged Arc1-12KR was purified from the supernatant by sequential chromatography using HiTrap Q, Resource Q anion exchange columns, and finally the Superdex 200 Increase gel filtration column. Homogeneity of the protein preparation was confirmed by performing sodium dodecyl sulfate-polyacrylamide gel electrophoresis and subsequently staining the gel with Coomassie Brilliant Blue. Protein concentrations were determined by measuring absorbance at 280 nm using the method of Pace et al.^46^, a refinement of the procedure originally described by Gill and von Hippel^47^.

### Physicochemical analyses

Analytical gel filtration chromatography was conducted using the Superdex 200 Increase 10/300 GL column (Cytiva, Tokyo, Japan) equilibrated with 20 mM Tris-HCl buffer (pH 8.0) containing 100 mM NaCl and 1 mM EDTA. Protein samples (0.1 mL) were applied to the column and eluted at 25 °C using a flow rate of 0.7 mL/min. Elution profiles were monitored by absorbance at 280 nm. Apparent molecular masses were estimated from elution volumes by referencing a calibration curve generated with protein standards.

Far-UV CD spectra were recorded using a J-1100 spectropolarimeter (Jasco, Hachioji, Japan) equipped with a programmable temperature control unit. Protein samples were prepared at a concentration of 20 μM in 20 mM Tris-HCl buffer (pH 8.0) containing 100 mM NaCl and 1 mM EDTA. Spectra were acquired from 200 to 250 nm using a 0.1-cm path length quartz cuvette at 25 °C.

### Enzyme assays

Canonical NDK activity was assessed at 60 °C by measuring the transfer of the γ-phosphate group from GTP to ADP, resulting in the production of GDP and ATP. The assay buffer consisted of 50 mM HEPES (pH 8.0), 25 mM KCl, 10 mM (NH₄)₂SO₄, 2.0 mM (CH₃COO)₂Mg, 1.0 mM DTT, 5 mM ADP, and 5 mM GTP. Disproportionation activity of ADP was similarly quantified at 60 °C by monitoring ATP production, using an assay buffer containing 50 mM HEPES (pH 8.0), 25 mM KCl, 10 mM (NH₄)₂SO₄, 2.0 mM (CH₃COO)₂Mg, 1.0 mM DTT, and 5 mM ADP. In both cases, the amount of ATP produced was quantified using the luminescent Kinase-Glo Plus assay kit (Promega Japan, Tokyo, Japan), as described previously^48^. One unit of enzyme activity was defined as the amount of protein required to generate 1 μmol of ATP per minute under the assay conditions. To examine the dependence of ATP formation on ADP concentration, the reaction was performed at 60 °C using ADP as the sole nucleotide substrate. ATP production was monitored using the same assay buffer with varying concentrations of ADP from 0.3 to 5 mM. Initial velocities were determined from the linear phase of ATP formation.

### HPLC analysis

The ability of Arc1-12KR to catalyze the disproportionation of ADP into ATP and AMP was confirmed by HPLC analysis. The reaction mixture contained 50 mM HEPES (pH 8.0), 25 mM KCl, 10 mM (NH₄)₂SO₄, 2.0 mM (CH₃COO)₂Mg, 1.0 mM DTT, and 5 mM ADP. Immediately after the addition of Arc1-12KR, the reaction solution was passed through a Millipore filter unit (0.45 μm pore size) and injected into the HPLC system (e-HPLC Kotori type A2, Uniflows, Tokyo, Japan) equipped with a PEGASIL ODS SP100 AQ column (Φ6×30 mm, Senshu Kagaku, Tokyo, Japan). Bound material was subsequently eluted from the column with 200 mM potassium phosphate (pH 6.0) at a flow rate of 1 mL/min. Nucleotides were detected by UV absorbance at 265 nm. A second sample, in which the Arc1-12KR reaction was incubated at 60 °C for 10 min, was also analyzed in the same manner. For reference, a chromatogram of standard solution of ATP and AMP mixture was also obtained (Fig. S3).

To assess the reverse reaction, an assay buffer containing 50 mM HEPES (pH 8.0), 25 mM KCl, 10 mM (NH₄)₂SO₄, either 2 or 10 mM (CH₃COO)₂Mg, 1.0 mM DTT, plus 5 mM each of AMP and ATP was used. In addition, a reaction mixture containing 5 mM GDP as a potential substrate was analyzed under the same experimental conditions used for the ADP disproportionation assay.

### X-ray crystallography

To obtain crystals of the simplified NDK Arc1-12KR, 200 nL of protein solution (15.4 mg/mL), supplemented with 5 mM ADP, was mixed with an equal volume of reservoir solution consisting of 20% (w/v) PEG3350 and 100 mM CHES-NaOH buffer (pH 9.5), using the sitting-drop vapor diffusion method. Crystallization drops were incubated at 20 °C. Crystals were harvested and briefly soaked in cryoprotectant solution containing 20% (w/v) PEG3350, 100 mM CHES-NaOH (pH 9.5), and 20% glycerol, followed by flash-cooling in liquid nitrogen. X-ray diffraction data were collected at beamline BL32XU of SPring-8 using the fully automated ZOO system^49–52^. Multiple small-wedge datasets (30° per wedge) were acquired from single crystals and automatically processed and merged using KAMO with the XDS suite^53, 54^.

Although ADP was included in the crystallization conditions to potentially stabilize the active site conformation, no clear electron density corresponding to ADP was observed in the final structure. The crystal structure was solved by molecular replacement with Phaser (v2.8.2) in the PHENIX suite^55^, using a ColabFold-predicted structure (v1.3.0)^56^ as the search model. Manual model building was performed in Coot (v0.8.9.1)^57^, followed by iterative refinement with phenix.refine (v1.14_3260) and validation with MolProbity. The final model has been deposited in the Protein Data Bank under accession code 9XEJ. Crystallographic data collection and refinement statistics are summarized in Table S1.

### Molecular docking and MD simulations

The ligand structure of ADP was built and energy-minimized using Avogadro^58^, followed by further optimization with the B3LYP/def2-SVP level of theory in ORCA 6.0^59^ to obtain accurate geometries and atomic charges. The crystal structure of Arc1-12KR was used as the receptor model, and the ligand position was initially guided by superimposing the Mg^2+^-bound NDK structure (PDB ID: 2DY9). Molecular docking was performed using an integrative modeling approach with HADDOCK 2.4^60^. This procedure included rigid-body, semi-flexible, and final flexible refinement steps. The resulting complexes were scored using HADDOCK’s integrated scoring function, and the top-ranked model was selected for MD simulations.

MD simulations were conducted for both the apo and ADP-bound forms of Arc1-12KR using GROMACS 2024.3^61^ with the CHARMM36^62^ force field. Each system was solvated in a TIP3P water box, neutralized with Na⁺ and Cl⁻ ions, and energy-minimized using the steepest descent method. Equilibration was performed for 5 ns each under NVT and NPT ensembles at 310 K and 1 bar, followed by a 1 μs production run without restraints. Bond constraints were applied using the LINCS algorithm^63^, and long-range electrostatics were treated with the PME method^64^ (1.2 nm cutoff). Simulations were repeated to ensure reproducibility.

Hydrogen-bonding interactions and water occupancy near the ligand-binding pocket were analyzed over the trajectory. Water molecules were considered conserved if they maintained interactions for more than 90 % of the simulation frames^65–67^. Spatial clustering of the water molecules was visualized using PyMOL (http://www.pymol.org) and VMD^68^.

### Synthesis of Arc1-12KR mutant genes

The genes for four single mutants (R85M, R102M, D115A, D118A) and two double mutants (R85M/R102M, D115A/D118A) were artificially synthesized by Eurofins Genomics. NdeI and BamHI restriction recognition sites were added upstream and downstream of the synthesized genes, respectively. The synthetic genes were digested with NdeI and BamHI (New England Biolabs, Tokyo, Japan) and ligated into the corresponding restriction recognition sites of plasmid pET15b. Expression, purification, and subsequent enzymatic analysis of each mutant protein, carrying an N-terminal His₆-tag, were performed as described above.

## Supporting information

DataS2

DataS3

## Acknowledgments

This work was supported by JSPS KAKENHI (Grant numbers 19K21903, 21H01200 and 23K20903) to SA; and by the Astrobiology Center Program of the National Institutes of Natural Sciences (Grant Numbers AB022003, AB031007 and AB0609) to SA.

## Author Contributions

S.A. designed research; S.Y., S.D., and S.A. performed research; S.Y., S.D., and S.A. analyzed data and wrote the manuscript; and all authors discussed the results and reviewed the final manuscript.

## Data Availability

The atomic coordinate file of Arc1-12KR is deposited in PDB under the accession code 9XEJ. Experimental data of biochemical characterization and the sequences of all constructs used in this study can be found in Supplementary Information. Requests for materials should be directed to the corresponding author.

