## Supplementary Information for "Restricted amino acid diversity alters enzymatic phosphoryl-transfer catalysis"

### Emergence of a potentially ancestral ATP-synthesizing activity under prebiotic amino acid constraints

**This PDF file includes:**

Figures S1 to S11
Tables S1
Data S1
Legends for Data S2 and Data S3

**Other supplementary materials for this manuscript include the following:**

Data S2, Data S3

```
             2                                                         60
Arc1         ERTFVMIKPDGVQRGLIGEIISRFERKGLKIVAMKMMRISREMAEKHYAEHREKPFFSA
Arc1-16      ERTFVMIKPDGVQRGLIGEIISRFERKGLKIVAMKMMRISREMAEKLLAELREKPFFSA
Arc1-13FKR   ERTFVLIKPDGVARGLIGEIISRFERKGLKIVALKLLRISRELAEKLLAELREKPFFSA
Arc1-13KMR   ERTLVMIKPDGVARGLIGEIISRLERKGLKIVAMKMMRISREMAEKLLAELREKPLLSA
Arc1-12KR    ERTLVLIKPDGVARGLIGEIISRLERKGLKIVALKLLRISRELAEKLLAELREKPLLSA
Arc1-11K     EKTLVLIKPDGVAKGLIGEIISKLEKKGLKIVALKLLKISKELAEKLLAELKEKPLLSA
Arc1-11R     ERTLVLIRPDGVARGLIGEIISRLERRGLRIVALRLLRISRELAERLLAELRERPLLSA
Arc1-10      EETLVLIEPDGVAEGLIGEIISELEEEGLEIVALELLEISEELAEELLAELEEEPLLSA


            61                                                         120
Arc1        LVDYITSGPVVAMVLEGKNAVEVVRKMVGATNPKEAAPGTIRGDFGLDVGKNVIHASDSP
Arc1-16     LVDLITSGPVVAMVLEGKDAVEVVRKMVGATDPKEAAPGTIRGDFGLDVGKLVIDASDSP
Arc1-13FKR  LVDLITSGPVVALVLEGKDAVEVVRKLVGATDPKEAAPGTIRGDFGLDVGKLVIDASDSP
Arc1-13KMR  LVDLITSGPVVAMVLEGKDAVEVVRKMVGATDPKEAAPGTIRGDLGLDVGKLVIDASDSP
Arc1-12KR   LVDLITSGPVVALVLEGKDAVEVVRKLVGATDPKEAAPGTIRGDLGLDVGKLVIDASDSP
Arc1-11K    LVDLITSGPVVALVLEGKDAVEVVKKLVGATDPKEAAPGTIKGDLGLDVGKLVIDASDSP
Arc1-11R    LVDLITSGPVVALVLEGRDAVEVVRRLVGATDPREAAPGTIRGDLGLDVGRLVIDASDSP
Arc1-10     LVDLITSGPVVALVLEGEDAVEVVEELVGATDPEEAAPGTIEGDLGLDVGELVIDASDSP


            121                139
Arc1        ESAEREISLFFKDEELVEW
Arc1-16     ESAEREISLFFKDEELVEW
Arc1-13FKR  ESAEREISLFFKDEELVER
Arc1-13KMR  ESAEREISLLLKDEELVER
Arc1-12KR   ESAEREISLLLKDEELVER
Arc1-11K    ESAEKEISLLLKDEELVEK
Arc1-11R    ESAEREISLLLRDEELVER
Arc1-10     ESAEEEISLLLEDEELVEE
```

**Figure S1.** A multiple amino acid sequence alignment of Arc1 and its reduced amino acid set variants. N-terminal residues were omitted from this alignment. Residue numbers are shown above the sequences. Active-site residues of Arc1 are highlighted in cyan, while residues subjected to site-directed mutagenesis in the Arc1-12KR variant are highlighted in green.

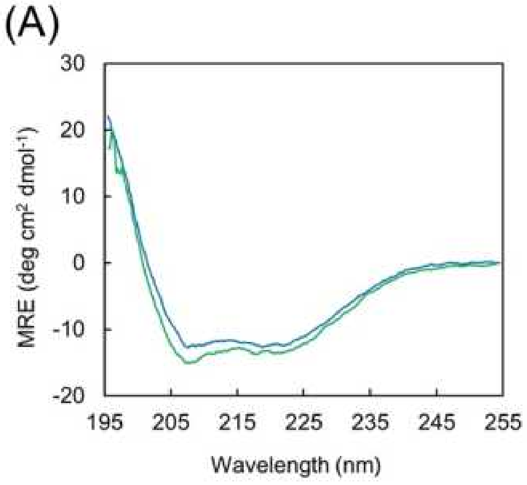


**Figure S2.** Comparison between untagged (green) and N-terminal His$_6$-tagged (blue) Arc1-12KRs. **(A)** Far-UV CD spectra. The proteins were analyzed at a concentration of 20 µM in 20 mM Tris-HCl, pH 8.0, 100 mM NaCl, and 1 mM EDTA. Spectra were obtained using a 0.1-cm path length quartz cell at 25 °C. **(B)** Time course of ATP production per µmol of protein from ADP as the sole substrate at 60°C. The assay solution was composed of 50 mM HEPES (pH 8.0), 25 mM KCl, 10 mM $(NH_4)_2SO_4$, 2.0 mM $(CH_3COO)_2Mg$, 1.0 mM DTT, and 5 mM ADP. Each value is the average of three measurements.

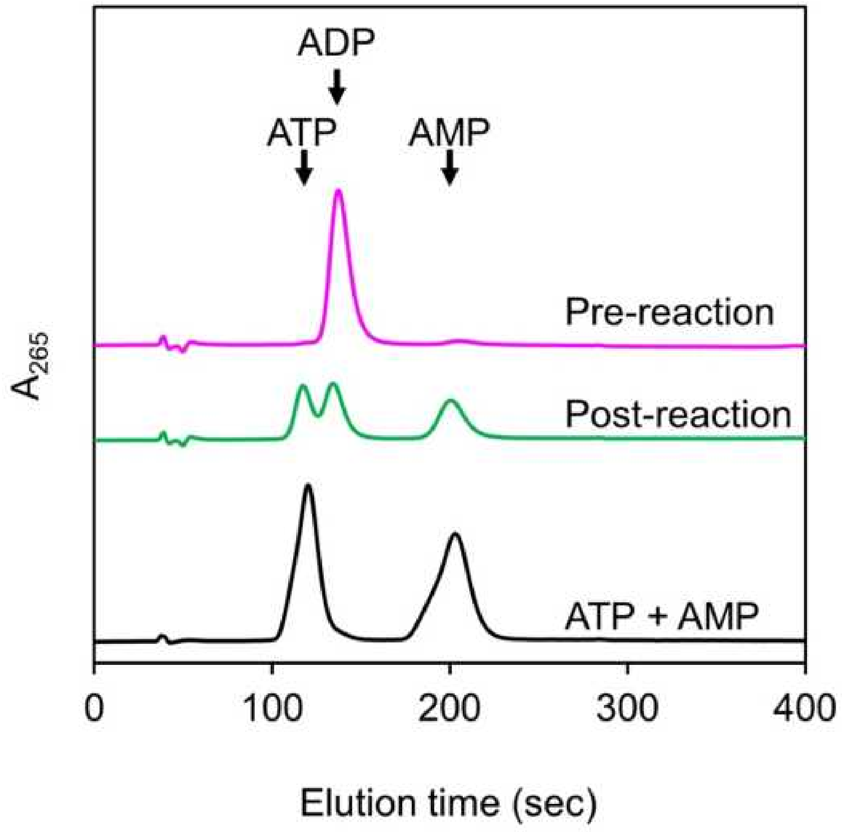


**Figure S3.** HPLC analysis of reaction mixtures using ADP as the sole substrate for Arc1-12KR. The chromatogram of the reaction mixture before enzyme addition is shown in magenta. The chromatogram obtained after incubation with 0.5 μM Arc1-12KR at 60°C for 10 min is shown in green. For comparison, the chromatogram of a standard mixture of ATP and AMP is shown in black.

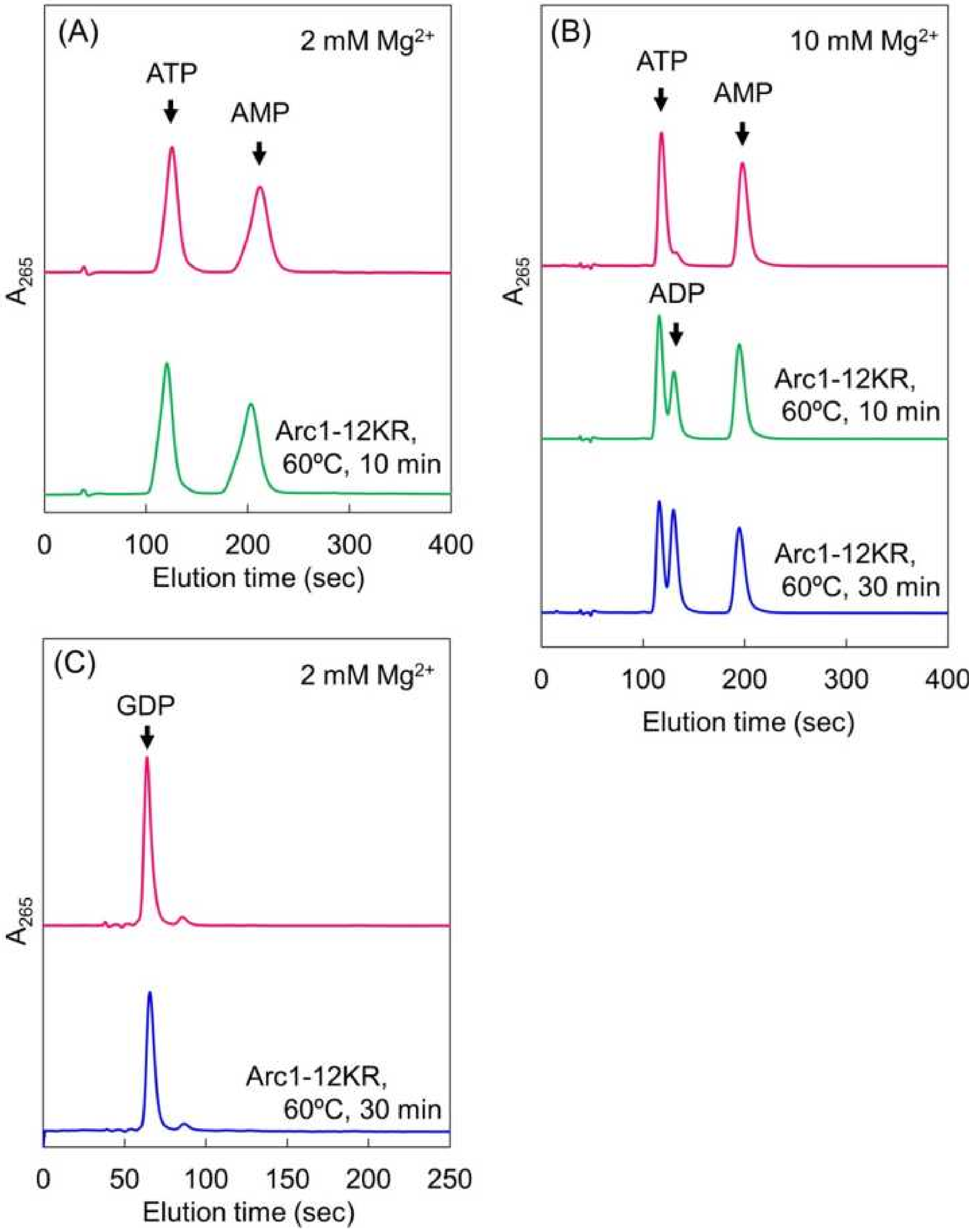


**Figure S4.** HPLC analysis of reaction mixtures using ATP and ADP (A, B) or GTP (C) as the substrate for Arc1-12KR. The $Mg^{2+}$ concentration was 2 mM in panels A and C and 10 mM in panel B. The chromatogram of the reaction mixture before enzyme addition is shown in magenta. Chromatograms obtained after incubation with 0.5 µM Arc1-12KR at 60°C for 10 min and 30 min are shown in green and blue, respectively.

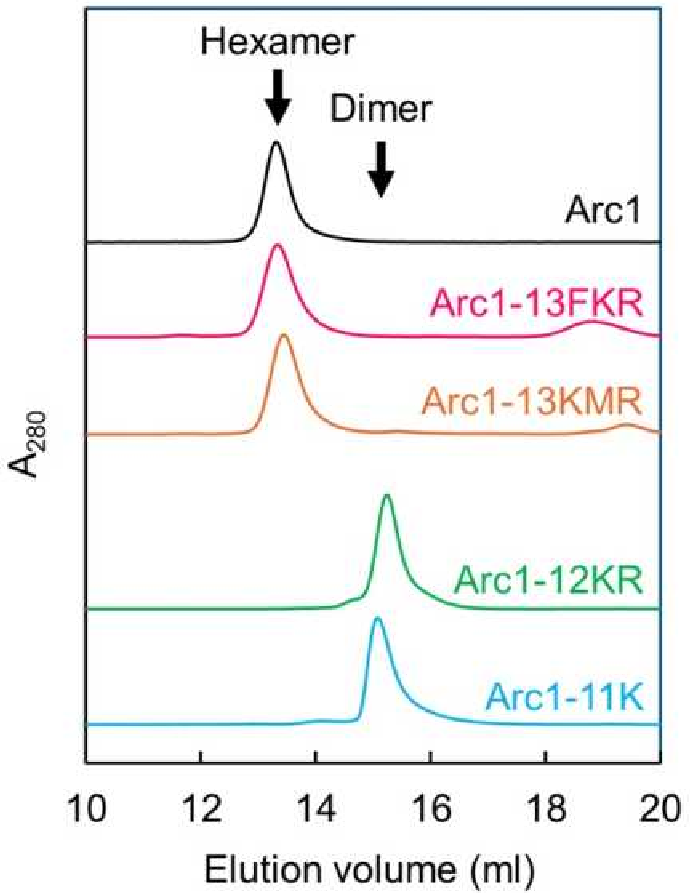


**Figure S5.** Analytical gel filtration chromatograms of Arc1 and its reduced amino acid set variants using a Superdex200 Increase column. Protein was applied at an initial concentration of 20 µM in 20 mM Tris-HCl, pH 8.0, 100 mM NaCl, and 1 mM EDTA with a flow rate of 0.7 mL/min. A280, absorbance at 280 nm.

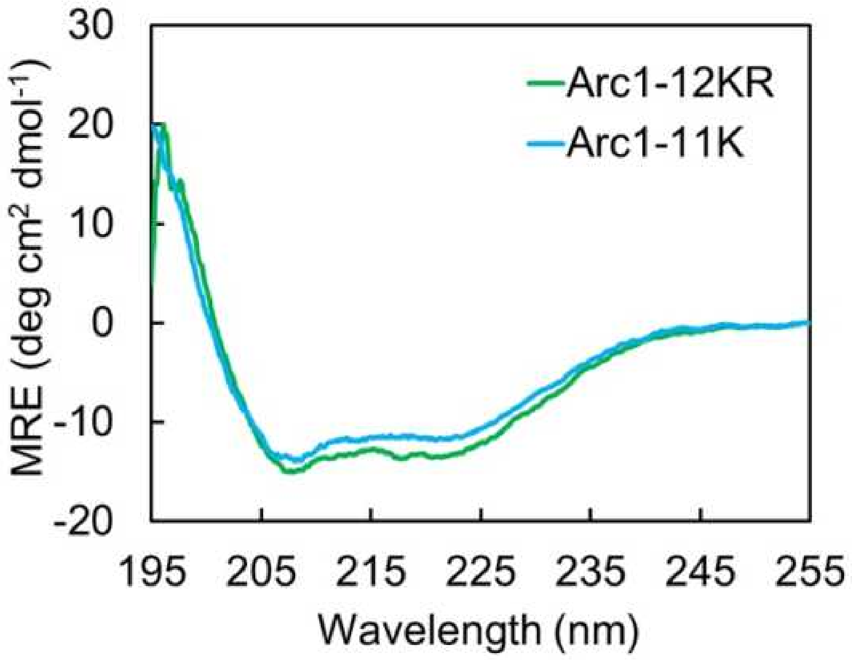


**Figure S6.** Far-UV CD spectra of Arc1-12KR and Arc1-11K. The proteins were analyzed at a concentration of 20 μM in 20 mM Tris-HCl, pH 8.0, 100 mM NaCl, and 1 mM EDTA. Spectra were obtained using a 0.1-cm path length quartz cell at 25 °C.

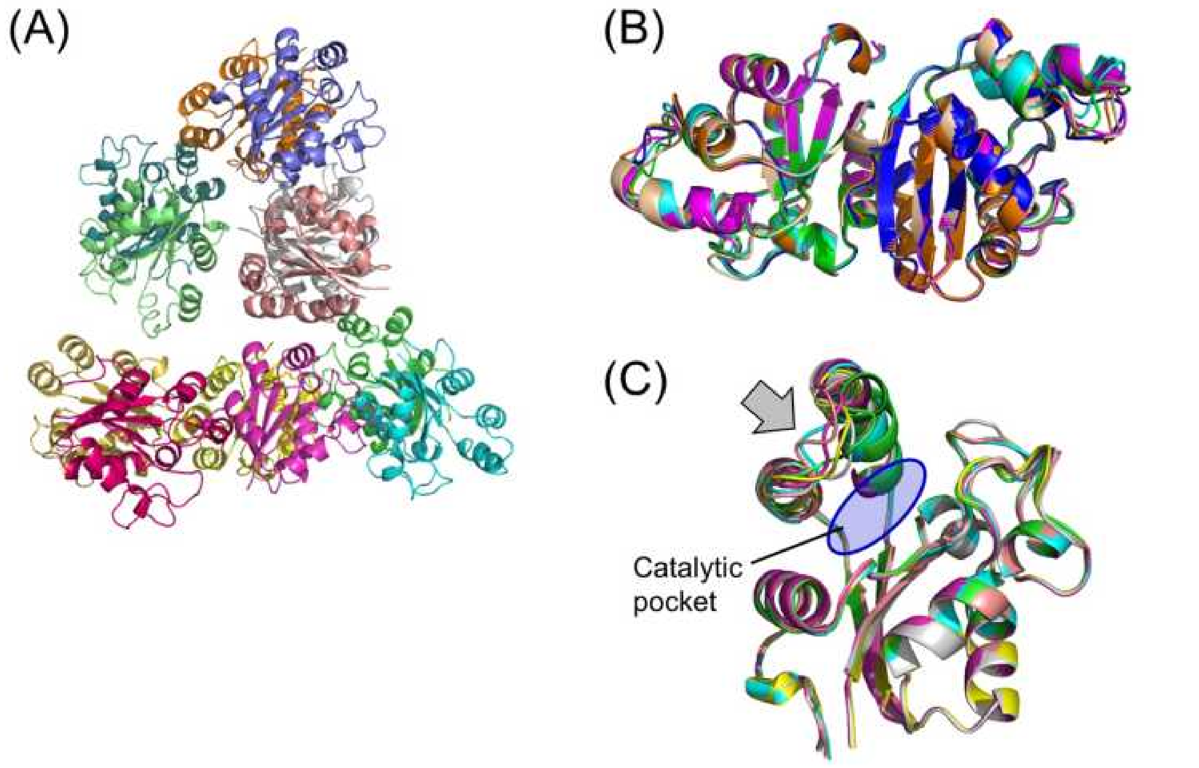


**Figure S7.** PyMOL (http://www.pymol.org) representation of the crystal structure of Arc1-12KR. **(A)** The six Arc1-12KR dimers present in the asymmetric unit are shown. For clarity, each polypeptide chain is rendered in a different color. **(B)** Superposition of all six dimers in the asymmetric unit. **(C)** Superposition of the monomeric subunits extracted from the six dimers shown in panel (B). The catalytic pocket is highlighted. A notable deviation is observed in the loop region (residues 49–59), as marked by an arrow.

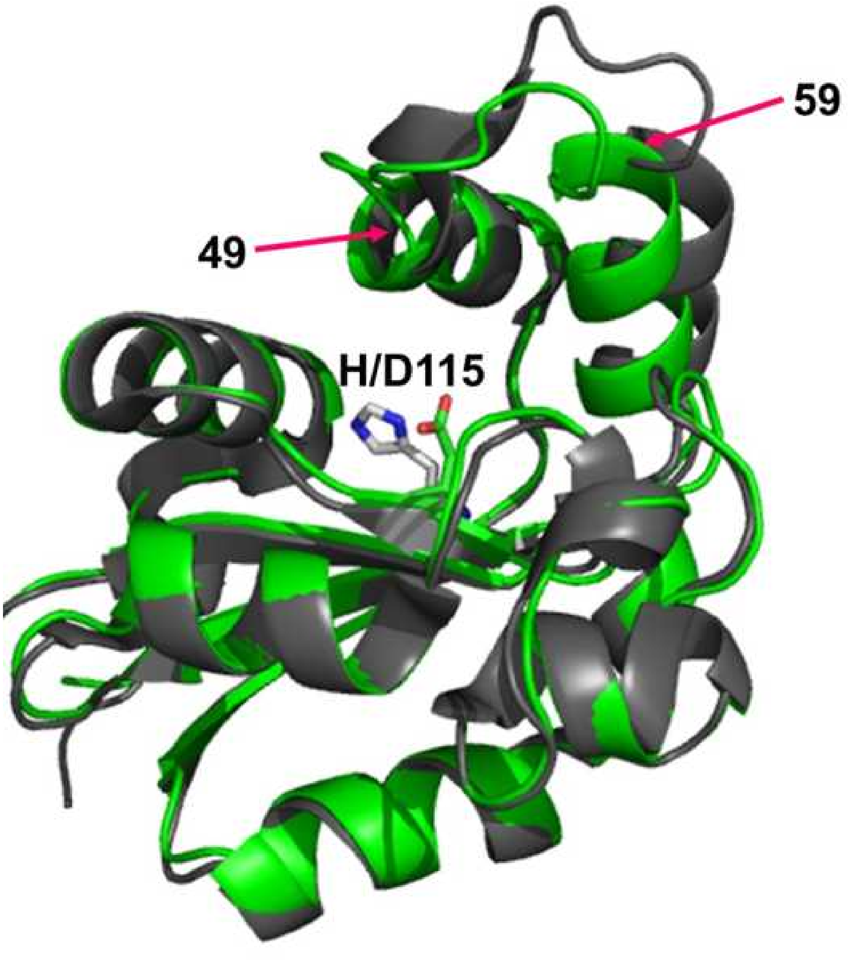


**Figure S8.** Superposition of the crystal structures of Arc1 (gray; PDB ID: 1VVT) and Arc1-12KR (green, PDB ID: 9XEJ). The side chains of the catalytically important residues at position 115 (His in Arc1; Asp in Arc1-12KR) are shown. A notable deviation is observed in the loop region comprising residues 49–59.

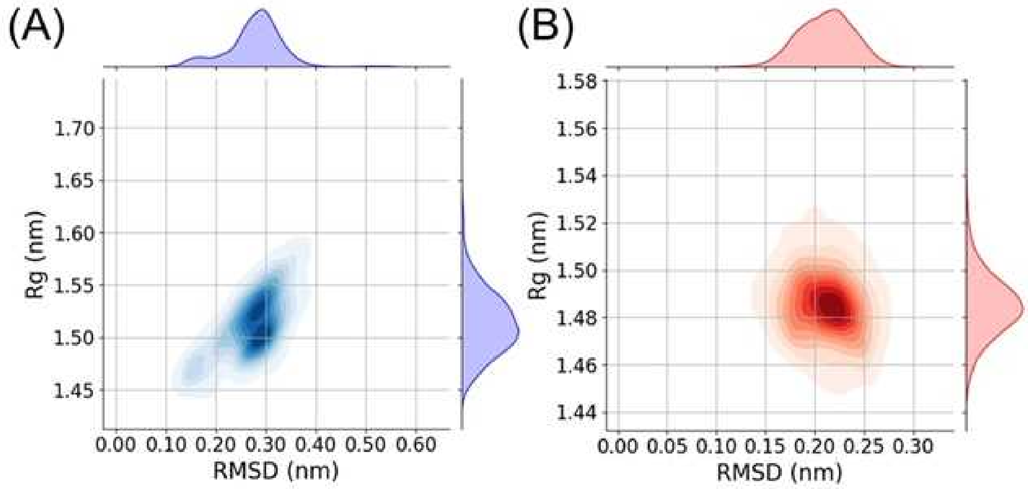


**Figure S9.** Root mean square deviation (RMSD) versus radius of gyration (Rg) plots of Arc1-12KR during molecular dynamics simulations. (A) Apo-Arc1-12KR. (B) ADP-bound Arc1-12KR (holo form). For the apo form, RMSD values ranged from approximately 0.25 to 0.37 nm, indicating relatively large structural fluctuations. In contrast, the holo form exhibited lower RMSD values, between about 0.15 and 0.26 nm, reflecting increased structural rigidity upon ADP binding. Consistent with this trend, the apo form showed Rg values ranging from ~1.50 to 1.55 nm, suggesting a slightly more extended and flexible conformation, whereas the holo form displayed Rg values between ~1.46 and 1.51 nm, indicative of a more compact structure.

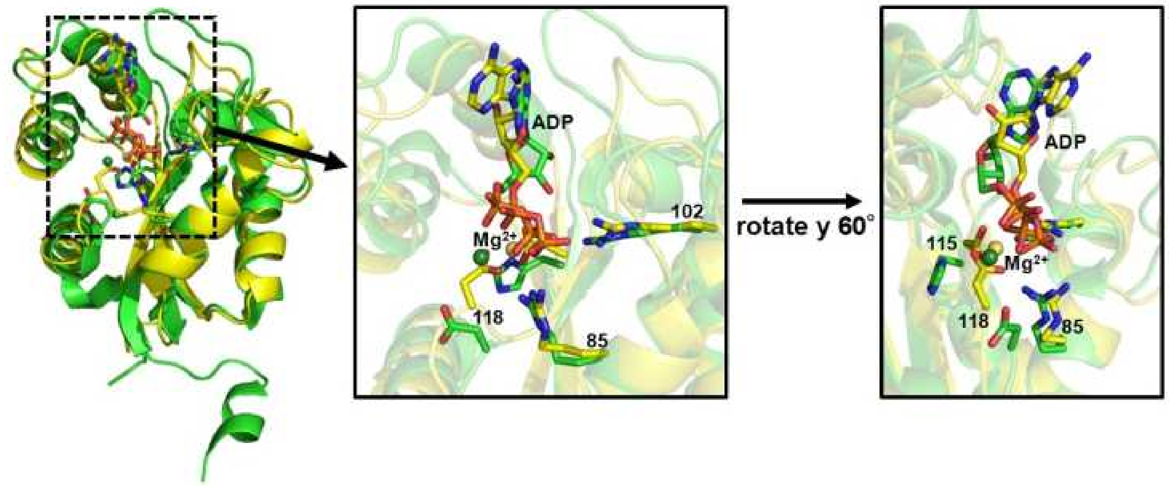


**Figure S10.** Superposition of the Arc1-12KR-ADP-$Mg^{2+}$ complex model (yellow) and the crystal structure of *P. horikoshii* NDK with bound ADP and $Mg^{2+}$ (green). The side chains of the four key residues (Arg85, Arg102, Asp/His115 and Asp118) are also shown.

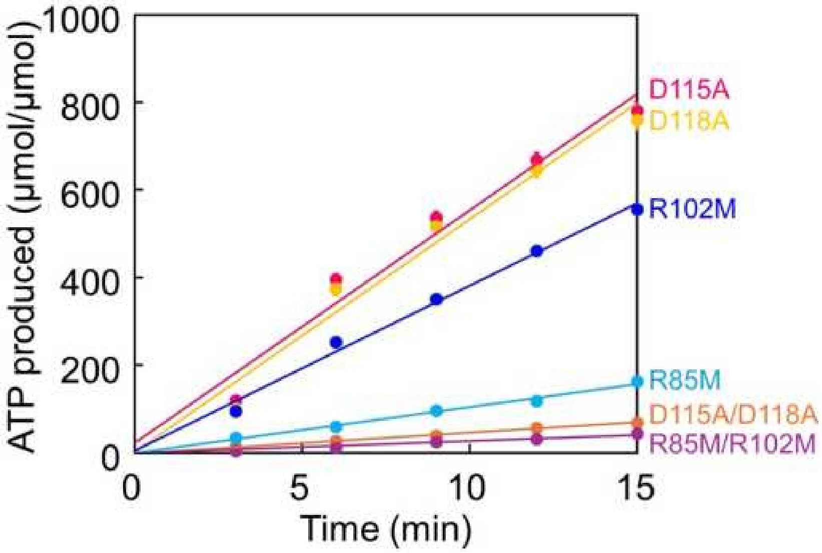


**Figure S11.** Time-course of ATP production per μmol of protein from ADP as the sole substrate by Arc12-related mutants. The assay solution was composed of 50 mM HEPES (pH 8.0), 25 mM KCl, 10 mM $(NH_4)_2SO_4$, 2.0 mM $(CH_3COO)_2Mg$, 1.0 mM DTT, and 5 mM ADP. Each value is the average of three independent measurements.

Table S1. Crystallographic data collection and refinement statistics.

| PDB ID | 9XEJ |
|---|---|
| Data collections | |
| Space group | p12(1)1 |
| Unit cell parameters | |
| a, b, c (Å) | 56,114.7, 128.8 |
| a, b, g (°) | 90.0, 95.0, 90.0 |
| Resolution (Å) | 46–3.34 (3.54–3.34) |
| Rsymm (%) | 25.5 (119.9) |
| Average I/sigma (I) | 9.00 (1.98) |
| Completeness (%) | 99.9 (100.1) |
| Redundancy | 9.0 (9.1) |
| Refinement | |
| Resolution (Å) | 45.53 – 3.34 |
| Unique reflections | 23 636 (2 369) |
| Rwork | 0.232 (0.2979) |
| Rfree | 0.258 (0.3116) |
| Number of atoms | 12492 |
| Protein atoms | 12492 |
| Ligands/ion | 0 |
| Water | 0 |
| R.m.s. deviations | |
| Bond lengths (Å) | 0.011 |
| Bond angles (°) | 1.54 |
| Ramachandran plot (%) | |
| Most favorable | 94.49 |
| Allowed | 5.51 |
| Disallowed | 0 |

**Data S1. Amino acid sequences of ancestral and *E. coli* IPMDHs in FASTA format.**

```
>Arc1
MERTFVMIKPDGVQRGLIGEIISRFERKGLKIVAMKMMRISREMAEKHYAEHREKPFFSALVDYITSG
PVVAMVLEGKNAVEVVRKMVGATNPKEAAPGTIRGDFGLDVGKNVIHASDSPESAEREISLFFKDEEL
VEW

>Arc1-16
MERTFVMIKPDGVQRGLIGEIISRFERKGLKIVAMKMMRISREMAEKLLAELREKPFFSALVDLITSG
PVVAMVLEGKDAVEVVRKMVGATDPKEAAPGTIRGDFGLDVGKLVIDASDSPESAEREISLFFKDEEL
VEW

>Arc1-13FKR
MERTFVLIKPDGVARGLIGEIISRFERKGLKIVALKLLRISRELAEKLLAELREKPFFSALVDLITSG
PVVALVLEGKDAVEVVRKLVGATDPKEAAPGTIRGDFGLDVGKLVIDASDSPESAEREISLFFKDEEL
VER

>Arc1-13KMR
MERTLVMIKPDGVARGLIGEIISRLERKGLKIVAMKMMRISREMAEKLLAELREKPLLSALVDLITSG
PVVAMVLEGKDAVEVVRKMVGATDPKEAAPGTIRGDLGLDVGKLVIDASDSPESAEREISLLLKDEEL
VER

>Arc1-12
MERTLVLIKPDGVARGLIGEIISRLERKGLKIVALKLLRISRELAEKLLAELREKPLLSALVDLITSG
PVVALVLEGKDAVEVVRKLVGATDPKEAAPGTIRGDLGLDVGKLVIDASDSPESAEREISLLLKDEEL
VER

>Arc1-11K
MEKTLVLIKPDGVAKGLIGEIISKLEKKGLKIVALKLLKISKELAEKLLAELKEKPLLSALVDLITSG
PVVALVLEGKDAVEVVKKLVGATDPKEAAPGTIKGDLGLDVGKLVIDASDSPESAEKEISLLLKDEEL
VEK

>Arc1-11R
MERTLVLIRPDGVARGLIGEIISRLERRGLRIVALRLLRISRELAERLLAELRERPLLSALVDLITSG
PVVALVLEGRDAVEVVRRLVGATDPREAAPGTIRGDLGLDVGRLVIDASDSPESAEREISLLLRDEEL
VER

>Arc1-10
MEETLVLIEPDGVAEGLIGEIISELEEEGLEIVALELLEISEELAEELLAELEEEPLLSALVDLITSG
PVVALVLEGEDAVEVVEELVGATDPEEAAPGTIEGDLGLDVGELVIDASDSPESAEEEISLLLEDEEL
VEE
```

**Data S2 (separate file). Data of activity measurements for Arc1 and its reduced amino acid set variants in Excel format.** All measurements were performed in triplicate. Data are shown as the mean and standard error of ATP produced per µmol of enzyme at each reaction time point.

**Data S3 (separate file). The number of occurrences of each amino acid at every position in a multiple sequence alignment of 309 extant NDK sequences in Excel format.**
